# Astrocytes are implicated in BDNF-mediated enhancement of hippocampal long-term potentiation

**DOI:** 10.1101/2021.03.30.437538

**Authors:** Joana I. Gomes, João Jesus, Renata Macau, Joana Gonçalves-Ribeiro, Sara Pinto, Carolina Campos Pina, Adam Armada-Moreira, Ana Maria Sebastião, Sandra H. Vaz

## Abstract

It is known that astrocytes, by the Ca^2+^-dependent release of gliotransmitters, which then act in pre- and post-synaptic receptors, modulate neuronal transmission and plasticity. Thus, hippocampal θ-burst long-term potentiation (LTP), which is a form of synaptic plasticity, can be modulated by astrocytes, since these cells release gliotransmitters that are crucial for the maintenance of LTP. Therefore, in this study, we hypothesized that the facilitatory action of BDNF upon LTP would involve astrocytes. To address that possibility, fEPSP recordings were performed in CA3-CA1 area of hippocampal slices from three different experimental models: Wistar rats where astrocytic metabolism was selectively reduced by a gliotoxin, the DL-fluoricitric acid (FC), IP3R2^−/−^ mice, which lack IP3R2-mediated Ca^2+^-signaling in astrocytes and dn-SNARE transgenic mice, in which the SNARE-dependent release of gliotransmittersis impaired. For the three models we observed that the astrocytic impairment abolished the excitatory BDNF effect upon hippocampal LTP, only while inducing LTP with a mild θ-burst stimulation paradigm. The present data shows for the first time that astrocytes play an active role in the facilitatory action of BDNF upon LTP, depending on stimulation paradigm.

## Introduction

Astrocytes, the main glial cell type in central nervous system (CNS), are in close contact with neurons leading to the recognition of a bidirectional communication between both neurons and astrocytes at the synapse (Araque *et al.*, 1999). These cells play a crucial modulatory role upon synaptic function since they respond to neurotransmitters through intracellular Ca2+ transients, that induce the release of gliotransmiters, which modulate neuronal excitability and synaptic function (Kofuji *et al.*, 2021). Among other functions, astrocytes modulate long-term potentiation (LTP), the neurophysiological basis for learning and memory (Bliss *et al.*, 1993), through the release of ATP (Pascual *et al.*, 2005), glutamate (Perea and Araque, 2007) and D-serine (Henneberger *et al.*, 2010). CA1 hippocampal LTP can be triggered when postsynaptic activity and astrocytic Ca^2+^-dependent glutamate release occur at the same time (Perea *et al.*, 2007). The Ca^2+^-dependent release of D-serine from CA1 hippocampal astrocytes controls NMDAR-dependent plasticity in the excitatory synapses nearby (Henneberger *et al.*, 2010). The extracellular adenosine, derived from astrocytic ATP (Bal-Price et al., 2002; Lalo et al., 2014), regulates synaptic transmission and modulates LTP (Pascual *et al.*, 2005).

Brain-derived neurotrophic factor (BDNF) is a neurotrophin with essential functions in neuronal survival, differentiation, and synaptic plasticity(G. Leal *et al.*, 2017; Sebastião *et al.*, 2011) that mediates its effects through the activation of TrkB receptors. Different TrkB receptor isoforms are generated by alternative splicing, namely one full-length form of TrkB (TrkB-fl) and two truncated TrkB-t (e.g. Sebastião et al., 2011) isoforms. The enhancement of LTP by BDNF (Korte *et al.*, 1995; Figurov *et al.*, 1996; Patterson *et al.*, 1996; Fontinha *et al.*, 2008) is achieved through the activation of TrkB-fl receptors (Xu *et al.*, 2000; Minichiello *et al.*, 2002), since hippocampal LTP is strongly impaired in both BDNF (Gottschalk *et al.*, 1999; Korte *et al.*, 1995; Patterson *et al.*, 1996) and TrkB (Xu *et al.*, 2000; Minichiello *et al.*, 2002) knockout mice. Besides BDNF effects on LTP, it has also been shown that in cultured astrocytes, BDNF mediates rapid Ca^2+^ transient through TrkB-t receptor, that results from G protein-dependent PLC activation and release of Ca^2+^ from Ins(1,4,5)P3-sensitive stores (Rose *et al.*, 2003). In neuromuscular junction was shown that BDNF elicits a Ca^2+^ response in perisynaptic Schwann cells thought TrkB-t receptor (Todd *et al.*, 2007). Moreover, BDNF overexpression from astrocytes lead to an improvement of cognitive performance in an Alzheimer’s mice model (De Pins *et al.*, 2019). Besides astrocytic BDNF-mediated modulation of Ca^2+^ transients, this neurotrophic factor is able to induce the release of glutamate from astrocytes through a Ca^2+^ dependent mechanism (Pascual et al, 2001), which may contribute to the enhancement of LTP.

Although it is well established that both BDNF and astrocytes modulate LTP, nothing is known about the role that astrocytes may play upon BDNF regulation of LTP. Here, we have investigated the functional role of astrocytes upon the enhancement of hippocampal LTP by BDNF in three complementary approaches: i) under the pharmacological astrocytic blockade with the gliotoxin fluorocitrate (FC) (Swanson *et al.*, 1994); ii) in a transgenic mice that lacks astrocytic release of gliotransmitters (dn-SNARE mice) (Pascual *et al.*, 2005); and iii) in a transgenic mice that lacks astrocytic Ca^2+^ signalling (IP3R2^−/−^ mice) (Li *et al.*, 2005). By performing electrophysiological recordings, in these three considered approaches, we unveiled that astrocytes control the effect of BDNF upon synaptic plasticity in the hippocampus. We thus identified a new role of astrocytes in the CNS - to fine-tune synaptic plasticity mediated by neurotrophins.

## Methods

### Experimental models

Animals were maintained in controlled temperature (21 ± 1°C) and humidity (55 ± 10%) conditions with a 12:12 h light/dark cycle and access to food and water *ad libitum.* All procedures were carried out according to the European Union Guidelines for Animal Care (European Union Council Directive 2010/63/EU) and Portuguese law (DL 113/2013) with the approval of the Institutional Animal Care and Use Committee. Care was taken to minimize the number of animals sacrificed.

#### Generation of 1,4,5-triphosphate (IP3) type-2 receptor (R2) knockout (IP3R2^−/−^) mice

A mouse strain with targeted deletion of Itpr2 gene was kindly supplied for this project by Prof. João F. Oliveira (U. Minho, Portugal) (Guerra-Gomes et al., 2020), under agreement with Prof. Ju Chen (U.C. San Diego, USA) (Li et al., 2005), who have generate the IP3R2^−/−^ mice. IP3R2^−/−^ and their respective littermate WT controls (IP3R^+/+^) were obtained by mating couples of IP3R2^+/−^. Collected toe tissue was used for DNA extraction and subsequent genotyping by polymerase chain reaction (PCR) analysis using WT (Forward (F), 5’-ACCCTGATGAGGGAAGGTCT-3’; Reverse (R), 5’-ATCGATTCATAGGGCACACC-3’) and mutant allele specific primers (neo-specific primer: F, 5’-AATGGGCTGACCGCTTCCTCGT-3’; R, 5’-TCTGAGAGTGCCTGGCTTTT-3’) as previously described(Li *et al.*, 2005).

#### Generation of dn-SNARE mice

A dn-SNARE strain was kindly supplied for this project by Prof. João F. Oliveira (U. Minho, USA) (Sardinha VM et al., 2017), under agreement with Prof. Philip Haydon (Tufts U., USA) (Pascual *et al.*, 2005), and were maintained in the C57Bl6/J genetic background. The generation of dn-SNARE mice was performed as previously described (Pascual *et al.*, 2005). Briefly, the dn-SNARE mice and wild-type (WT) littermates were obtained by crossing two distinct transgenic mouse lines: hGFAP.tTA mice, where the expression of tetracycline transactivator (tTA) is mediated by the astrocyte-specific human glial fibrillary acidic protein (hGFAP) promoter and tetO.dnSNARE, in which the dominant-negative domain of vesicular SNARE VAMP2/synaptobrevin II, as well as, the reporter gene for enhanced green fluorescence protein (EGFP) are co-expressed under the control of the tetracycline operator (tetO). The generated animals will express Lac-Z, EGFP and SNARE transgenes on their astrocytes, but not on their neurons which leads to the selective impairment of vesicular gliotransmitters release from astrocytes. The conditional expression of the dnSNARE transgenes caused interference with the SNARE complex formation and consecutive blockade of exocytosis specifically in astrocytes (astrocytes derived from dnSNARE mice displayed a 91% reduction in the number of fusion events) (Sultan *et al.*, 2015), impairing the vesicular release of gliotransmitters. Doxycycline (Dox, Sigma-Aldrich, St. Louis, Missouri, EUA) administration in the drinking water (25 μL/mL), prevents the transgene expression, consequently leading to the inhibition of SNARE, Lac-Z and EGFP expression, resulting in the functional gliotransmission process. Animals genotype was confirmed by PCR analysis, where mice negative (wild-type) or positive for both transgenes (dn-SNARE) were tested, while mice expressing only single transgenes (GFAP-tTA or tetO.dnSNARE) were not included. Collected toe tissue was used for DNA extraction and subsequent genotyping by PCR analysis using primers to identify the tTA, tetO and HSF-1 genes: tTA-F, 5’-ACTCAGCGCTGTGGGGCATT-3’ and tTA-R, 5’-GGCTGTACGCGGACCCACTT-3’; tetO-F, 5’-TGGATAAAGAAGCTCATTAATTGTCA-3’ and tetO-R, 5’-GCGGATCCAGACATGATAAGA-3’; HSF-1-F, 5’-TCTCCTGTCCTGTGTGCCTAGC-3’ and HSF-1-R, 5’-CAGGTCAACTGCCTACACAGACC.

### Hippocampal slices preparation

Hippocampal slices were prepared as routinely in our lab (Rei *et al.*, 2020) from Wistar rats, IP3R2-KO and dn-SNARE mice with 8-12 weeks old. Wistar rats (Charles River, Barcelona, Spain) were deeply anesthetized with isoflurane (Esteve, Barcelona, Spain) and sacrificed by decapitation for hippocampal slice preparation. IP3R2-KO and dn-SNARE mice were sacrificed by decapitation after cervical displacement and the brain was rapidly removed in order to prepare hippocampal slices. The hippocampi were dissected in ice-cold artificial cerebrospinal fluid (aCSF) containing (in mM): 124 NaCl, 3 KCl, 1.2 NaH_2_PO_4_, 25 NaHCO_3_, 2 CaCl_2_, 1 MgSO_4_ and 10 glucose), which was continuously gassed with 95% O_2_ and 5% CO_2_. The hippocampal slices were quickly cut perpendicularly to the long axis of the hippocampus (400 μm thick) with a McIlwain tissue chopper and allowed to recover functionally and energetically for at least 1h in a resting chamber filled with continuously oxygenated aCSF, at room temperature (22–25°C), before being set up for electrophysiological recordings.

### Extracellular recordings of fEPSPs

Following the recovery period, slices were transferred to a recording chamber for submerged slices (1ml capacity plus 5 ml dead volume) and were constantly superfused at a flow rate of 3ml/mim with aCSF kept at 32°C, gased with 95% O_2_ / 5%CO_2_. Evoked field excitatory postsynaptic potentials (fEPSP) were recorded extracellularly by using a microelectrode (4 to 8 MΩ resistance) filled with aCSF solution placed in the stratum radiatum of the CA1 area. fEPSP data were acquired using an Axoclamp-2B amplifier (Axon Instrumnets, Foster City, CA). fEPSPs were evoked by stimulation through a concentric electrode to the Schaffer collateral fibres. Each individual stimulus consisted of a 0.1 ms rectangular pulse applied once every 20s, except otherwise indicated. Averages of six consecutive responses were continuously acquired, digitized with the WinLTP program (Anderson and Colloingridge 2001) and quantified as the slope of the initial phase of the averaged fEPSPs (and the amplitude of presynaptic fibre voley while performing input/output analysis). The stimulus intensity was adjusted at the beginning of the experiment to obtain a fEPSP slope that corresponds to about 50 % of the maximal fEPSP slope.

#### Input/output curves

For input/output curves (I/O), after obtaining a stable baseline under the standard stimulation conditions, the stimulus intensity was increased by 20μA every 4 minutes (60-320 μA). The I/O curves were plotted as the fEPSP slope against the stimulus intensity, as the presynaptic fibber volley (PSFV) amplitude against stimulus intensity, and as the fEPSP slope against PSFV amplitude, which provides a measure of synaptic efficiency. The maximal slope values were obtained by extrapolation upon nonlinear fitting of the I/O curve, an F-test being used to determine differences between the parameters.

#### Long-term potentiation induction

LTP was induced by a applying θ-burst stimulation, since this pattern of stimulation is considered to be closer to what occurs physiologically in the hippocampus during episodes of learning and memory in living animals (Albensi *et al.*, 2007). Two different θ-burst stimulation paradigms were used: one, named as strong θ-burst LTP-inducing paradigm, consisting of 1 train of 15 bursts (200 ms inter-burst interval), each burst being composed by 4 pulses delivered at 100 Hz [1 × (15×4)]; the other named week θ-burst LTP-inducing paradigm, consisting of 1 train of 3 bursts (200 ms inter-burst interval), each burst being composed by 3 pulses delivered at 100 Hz [1 × (3×3)]. The strong θ-burst stimulation was chosen since is well documented the exogenous BDNF effect on LTP for this particular stimulus (Fontinha *et al.*, 2008). The weak θ-burst stimulation paradigm was the same used by Pascual and collaborators (2005) to demonstrate the LTP impairment in dn-SNARE mice.

After obtaining a stable recording of the fEPSP slope, one of θ-burst of stimuli described were applied, and the stimulus paradigm was then resumed to pre-burst conditions up to the end of the recording period (60 min after burst stimulation). LTP magnitude was quantified as the % change in the average slope of the fEPSP taken from 50-60 minutes after the induction of LTP as compared with the average slope of the fEPSP measured during the 10 minutes before the induction of LTP. Since each slice allows recordings from two independent pathways, LTP was recorded in the first pathway, after 60 min of LTP induction, BDNF (20 ng/mL) was added to the superfusion solution. After at least 30 min of BDNF perfusion, LTP was induced in the second pathway. BDNF remained in the bath until the end of the experiment. I/O data was acquired in a new slice by stimulating the second pathway. In any case, fEPSPs were recorded under basal stimulation conditions (standard stimulus intensity and frequency) and stability of fEPSP slope values were guaranteed for more than 10min before changing any protocol parameter. One or two slices per animal were tested in each experimental day.

#### Drugs

BDNF was generously provided by Regeneron Pharmaceuticals (Tarrytown, NY), DL-fluorocitric acid (FCA) (200μM), 2-p-(2-Carboxyethyl) phenethylamino-5’-N-ethylcarboxamidoadenosine (CGS 21680, 30 nM), 2-(2-Furanyl)-7-(2-phenylethyl)-7H-pyrazolo[4,3-e][1,2,4]triazolo[1,5-c]pyrimidin-5-amine (SCH 58261, 50nM) were purchased from Sigma (St. Louis, MO). CGS 21680 and SCH 58261 were made up into a 5 mM stock solution in dimethylsulfoxide (DMSO). DL-flurocitrate was prepared as described previously(Paulsen *et al.*, 1987) and aliquots were made up into a 3.23 mM stock solution. BDNF was supplied in a 1.0 mg/mL stock solution in 150 mM NaCl, 10 mM sodium phosphate buffer, and 0.004% Tween 20. Aliquots of these stock solutions were kept frozen at −20°C until further use.

#### Statistical analysis

Data are expressed as means ± SEM from n slices. All statistical analysis was performed using GraphPad (San Diego, CA, USA) Prism software. Two-sample comparisons were made using t tests or by one-way ANOVA with treatment as the between-subject factor, followed by Sidak’s *post hoc* test for multiple comparisons when comparing multiple experimental groups. Values of p<0.05 were considered to account for statistically significant differences.

## Results

### Astrocytes do not control BDNF effect upon LTP induced by strong-θ-burst stimulation

To evaluate the role of astrocyte signaling for the effect of BDNF on synaptic plasticity, LTP was assessed in the CA1-CA3 area of the hippocampal slices of Wistar rats, IP3R2^−/−^ mice and dn-SNARE mice, and their respective littermate WT controls. The θ-burst stimulation of the Schaffer-collateral pathway led to an initial enhancement of the fEPSP slope, followed by a decrease of the slope towards a stabilization above the values recorded before θ-burst stimulation (baseline).

In the Wistar rats, the strong θ-burst stimulation protocol induced a LTP magnitude, quantified as the % change in the average slope of the fEPSP before and 50–60 min after θ-burst stimulation (see Methods), of 36.7±9.23% (n=5), while in the same slices but in the presence of BDNF (20ng/mL) the same induction paradigm induced a LTP magnitude of 80.4±8.95% (n=5). Thus, as previously reported in literature (Jerónimo-Santos *et al.*, 2014; Fontinha *et al.*, 2008), BDNF increased significantly the magnitude of LTP (Figure 1A,C; one-way ANOVA following Holm-Sidak’s *post hoc* test, **p=0.003, F (3, 17) = 22,04; n=5). To selectively reduce the astrocytic metabolism we used the fluorocitrate (FC), a reversible toxin that inhibits the enzyme aconitase causing a disruption of the tricarboxylic acid cycle and a decrease of the exported glutamine (Gln) (Berg-Johnsen *et al.*, 1993; Hassel *et al.*, 1994), by interfering with the glutamate–Gln cycle (Largo *et al.*, 1996; Paulsen *et al.*, 1987; Swanson *et al.*, 1994). In hippocampal slices of Wistar rats superfused with FC (200 μM) the same stimulation paradigm induced a decrease of fEPSP slope by −17.7±10.6% (n=5), corresponding to a marked and significant impairment of LTP when compared to hippocampal slices without FC superfusion (Figure 1B, C; one-way ANOVA following Holm-Sidak’s *post hoc* test θp=0.001, n=5). In Wistar rat hippocampal slices superfused with FC (200 μM) and BDNF (20ng/mL) the magnitude of LTP was 25.6±4.99%, which corresponds to a significant LTP potentiation by BDNF (Figure 1B, C one-way ANOVA following Holm-Sidak’s *post hoc* test **p=0.003, n=6).

**Figure 1.**
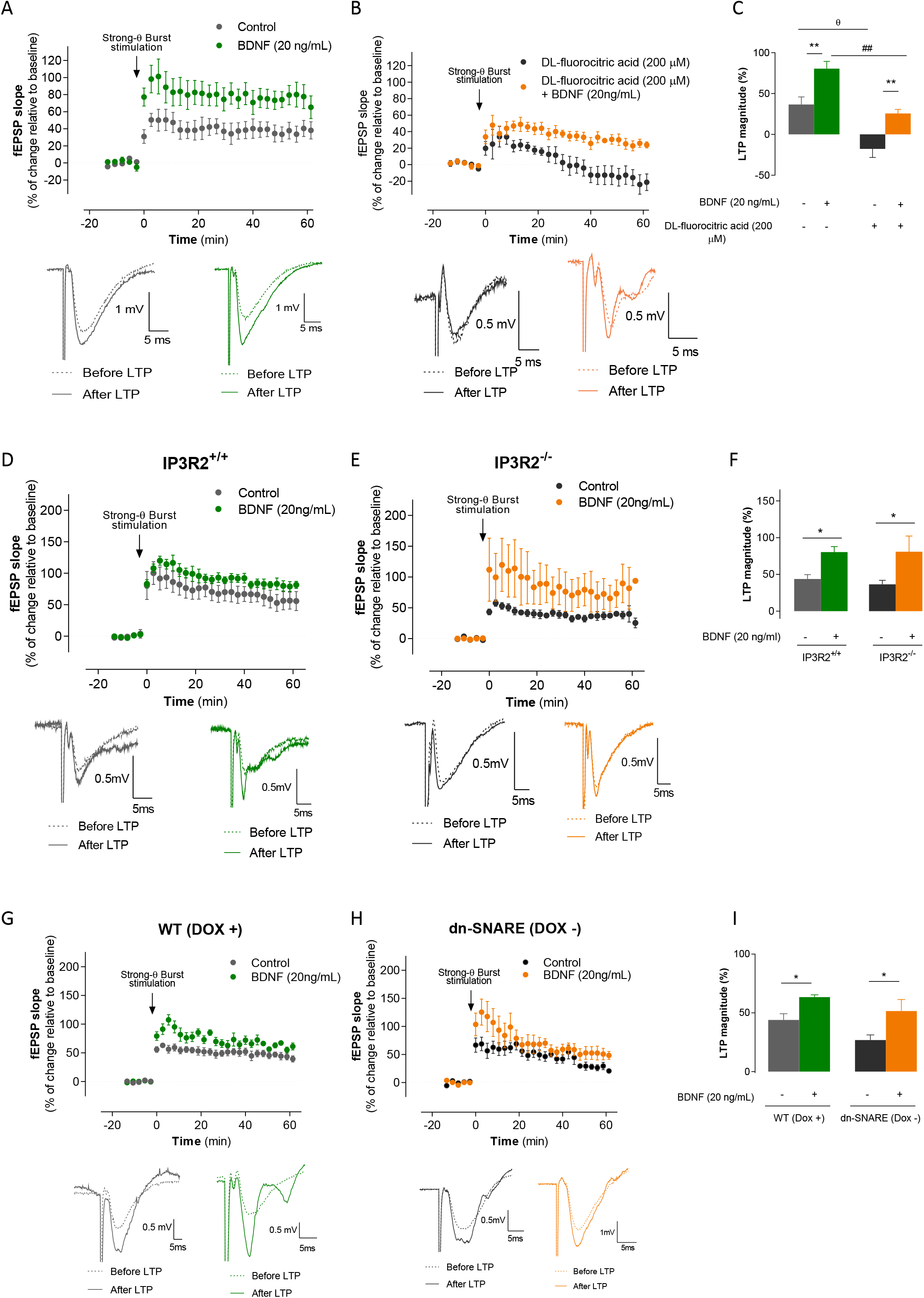
BDNF effect upon hippocampal LPT is not under astrocytic control while inducing LTP with a strong-θ-burst stimulation. In (A) is represented the average time course changes in the fEPSP slope induced by strong θ-burst stimulation in Wistar rats in the absence 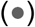 (n=5) or in the presence 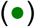 of BDNF (20 ng/mL) (n=5). In (B) is represented the average time course changes in the fEPSP slopes induced by strong θ-burst stimulation in Wistar rats in the presence of FC alone 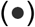 (n=5) or together with BDNF 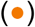 (20 ng/mL) (n=6). In (C) is represented the histogram depict LTP magnitude (change in fEPSP slope at 50-60 min) induced by strong θ-burst stimulation in Wistar rats in the presence or absence of FC and/or BDNF as indicated below each column. (D) represents the time course of averaged normalized changes in fEPSP slope after delivery (arrow) of a strong θ-burst to hippocampal slices from IP3R2^+/+^ in the absence 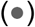 (n= 8) or in the presence 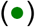 of BDNF (20 ng/mL) (n= 7). (E) Represents the time course of averaged normalized changes in fEPSP slope after delivery (arrow) of a strong θ-burst to hippocampal slices from IP3R2^−/−^ in the absence 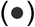 (n= 4) or in the presence 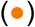 of BDNF (20 ng/mL) (n= 3). (F) corresponds to the histogram depict LTP magnitude (change in fEPSP slope at 50-60 min) induced by strong θ-burst stimulation in the absence or presence of BDNF (20 ng/mL) for each group of animals tested, as indicated below each column. (G) Represents the time course of averaged normalized changes in fEPSP slope after delivery (arrow) of a strong θ-burst to hippocampal slices from WT (Dox +) in the absence 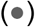 (n= 8) or in the presence 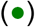 of BDNF (20 ng/mL) (n= 5). (H) Represents the time course of averaged normalized changes in fEPSP slope after delivery (arrow) of a strong θ-burst to hippocampal slices from dn-SNARE (Dox -) in the absence 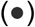 (n= 6) or in the presence 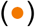 of BDNF (20 ng/mL) (n= 4). (I) corresponds to the histogram depict LTP magnitude (change in fEPSP slope at 50-60 min) induced by strong θ-burst stimulation in the absence or presence of BDNF (20 ng/mL) for each group of animals tested, as indicated below each column. The ordinates in (A), (B), (D), (E), (G) and (H) represent normalized fEPSP slopes where 0% corresponds to the averaged slope recorded for 10 min before θ-burst stimulation and the abscissa represents the time that average begun. Insets bellow each graph correspond to illustrative traces from representative experiments; each trace is the average of eight consecutive responses obtained for each group before (⋯) and 58-60 min after (-) θ-burst stimulation, and is composed of the stimulus artifact, followed by the presynaptic volley and the fEPSP. All values are presented as mean ± standard error of mean (SEM) from *n* independent observations. n.s (not significant): p> 0.05; *p≤ 0.05; **p ≤ 0.01; ^θ^p ≤ 0.01; ^##^p ≤ 0.001 (one-way ANOVA followed by Holm-Sidak’s *post hoc* test for multiple comparisons).

The role of astrocytes upon synaptic plasticity is dependent of intracellular Ca^2+^ signaling (Henneberger *et al.*, 2010) and gliotransmitter release (Pascual *et al.*, 2005). Since we observed that FC, due to its toxicity, completely impaired LTP, in the following experiments we used two different animal models that have impaired astrocytic Ca^2+^ signaling and gliotransmitter release, respectively, to further infer on the function of astrocytes for BDNF effect on synaptic plasticity. While performing fEPSPs recordings in IP3R2^−/−^ mice (Li *et al.*, 2005), which lacks IP3-mediated Ca^2+^ increases in astrocytes, and the respective littermate WT controls (IP3R2^+/+^), we observed that the strong-θ-burst stimulation applied in CA3-CA1 area of hippocampal slices from IP3R2^+/+^ mice induced a significant LTP magnitude increase of 43.7±5.9% (n=8) while in the same slices but in the presence of BDNF (20ng/mL) the same induction paradigm induced a LTP magnitude of 80.4±7.44% (n=7), corresponding to a significant LTP potentiation mediated by BDNF (Figure 1D, F; one-way ANOVA following Holm-Sidak’s *post hoc* test; *p=0.010, F (3, 18) = 6.914, n=7-8). Strong-θ-burst stimulation applied in hippocampal slices obtain from IP3R2^−/−^ mice, result in an increase of fEPSP slope of 36.4±5.6% (n=4) in the absence of BDNF (20ng/mL), while in its presence of 80.8±21.3% (n=3), which corresponds to a significant increase in LTP magnitude (Figure 1E, F; one-way ANOVA following Holm-Sidak’s *post hoc* test; *p=0.031, n=3-4).

In order to study the influence of gliotransmitters release on the BDNF effect upon hippocampal LTP we have used the dn-SNARE mice model which conditionally lacks astrocytic release of gliotransmitters by exocytosis. This conditional blockade of gliotransmitters release is achieved through the conditional expression of the dominant negative domain of vesicular SNARE protein synaptobrevin II (dn-SNARE), which interferes with the SNARE complex formation, impairing vesicular release (Sardinha *et al.*, 2017; Pascual *et al.*, 2005), being the expression of dn-SNARE transgene abolish by administration of doxycycline (DOX) in the drinking water of the animals. A strong θ-burst stimulation applied to hippocampal slices from WT (DOX+) mice result in an increase of fEPSP slope of 43.9±5.28% (n=8), whereas in the same slices but in the presence of BDNF (20ng/mL) the same induction paradigm enhanced the fEPSP slope to 63.3±1.99% (n=5), corresponding to a significant potentiation of LTP magnitude (Figure 1G, I; one-way ANOVA following Holm-Sidak’s *post hoc* test; *p=0.049; F (5, 24) = 5.79; n=5-8). For dn-SNARE (DOX+) animals, which release gliotransmitters in an exocytosis dependent mechanism, similar to WT animals, the magnitude of LTP for the same LTP induction protocol was 30.5±6.87% (n=5) in the absence of BDNF (20ng/mL), while in its presence was 5523±7.65% (n=3), that corresponds to a significant LTP potentiation (Supp. 1 A, C; unpaired t-test, *p=0.024, n=3-5). The obtained results in hippocampal slices from dn-SNARE (DOX-) mice, which lacks the release of gliotransmitters by exocytosis, show that the magnitude of the LTP, evoked by strong θ-burst stimulation, was 26.9±4.36% (n=6) in the absence of BDNF (20ng/mL), while in its presence was 51.5±9.87% (n=4), which corresponds to a significant LTP potentiation (Figure 1H, I; one-way ANOVA following Holm-Sidak’s *post hoc* test; *p=0.032, n=4-6). Altogether the results obtain from hippocampal slices of Wistar rats in presence of FC, IP3R2^−/−^ mice and dn-SNARE mice have shown that a strong θ-burst stimulation induces a significant LTP that is significantly potentiated by the presence of BDNF.

### Astrocytes contrinute to the BDNF effect upon LTP induced by mild-θ-burst stimulation

Since the ability of BDNF to facilitate LTP is more prominent when LTP is induced by a weaker than by a stronger θ-burst paradigm, next we explored if astrocytes could control BDNF effect upon hippocampal LTP while using a mild θ-burst stimulation to induce LTP. A mild-θ-burst stimulation applied in CA3-CA1 area of hippocampal slices from Wistar rats enhanced the fEPSP slope by 15.5±5.9% (n=7, Figure 2A, C). In the same slices but in the presence of BDNF (20ng/mL), mild-θ-burst stimulation increased fEPSP slope by 40.9±5.0% (n=6, Figure 2A, C) which corresponds to a significant LTP potentiation mediated by BDNF as previously reported (Fontinha *et al.*, 2008; Jerónimo-Santos *et al.*, 2014) (Figure 2A, C; one-way ANOVA following Holm-Sidak’s *post hoc* test; *p=0.024; F (5, 36) = 11.83; n=6-7). In hippocampal slices of Wistar rats superfused with FC (200 μM) the same mild stimulation paradigm induced a decrease of fEPSP slope by −11.7±6.70% (n=12), corresponding to a marked and significant impairment of LTP when compared to hippocampal slices without FC superfusion (Figure 2A, B, C; one-way ANOVA following Holm-Sidak’s *post hoc* test; θp=0.006, n=7-12). In slices superfused with FC, the presence of BDNF (20ng/mL) induced a LTP magnitude of −15.5±6.38% (n=7), corresponding to a significant loss of excitatory BDNF action on LTP in the presence of FC (Figure 2B, C; one-way ANOVA following Holm-Sidak’s *post hoc* test; ***p<0.001, n=6-7).

**Figure 2.**
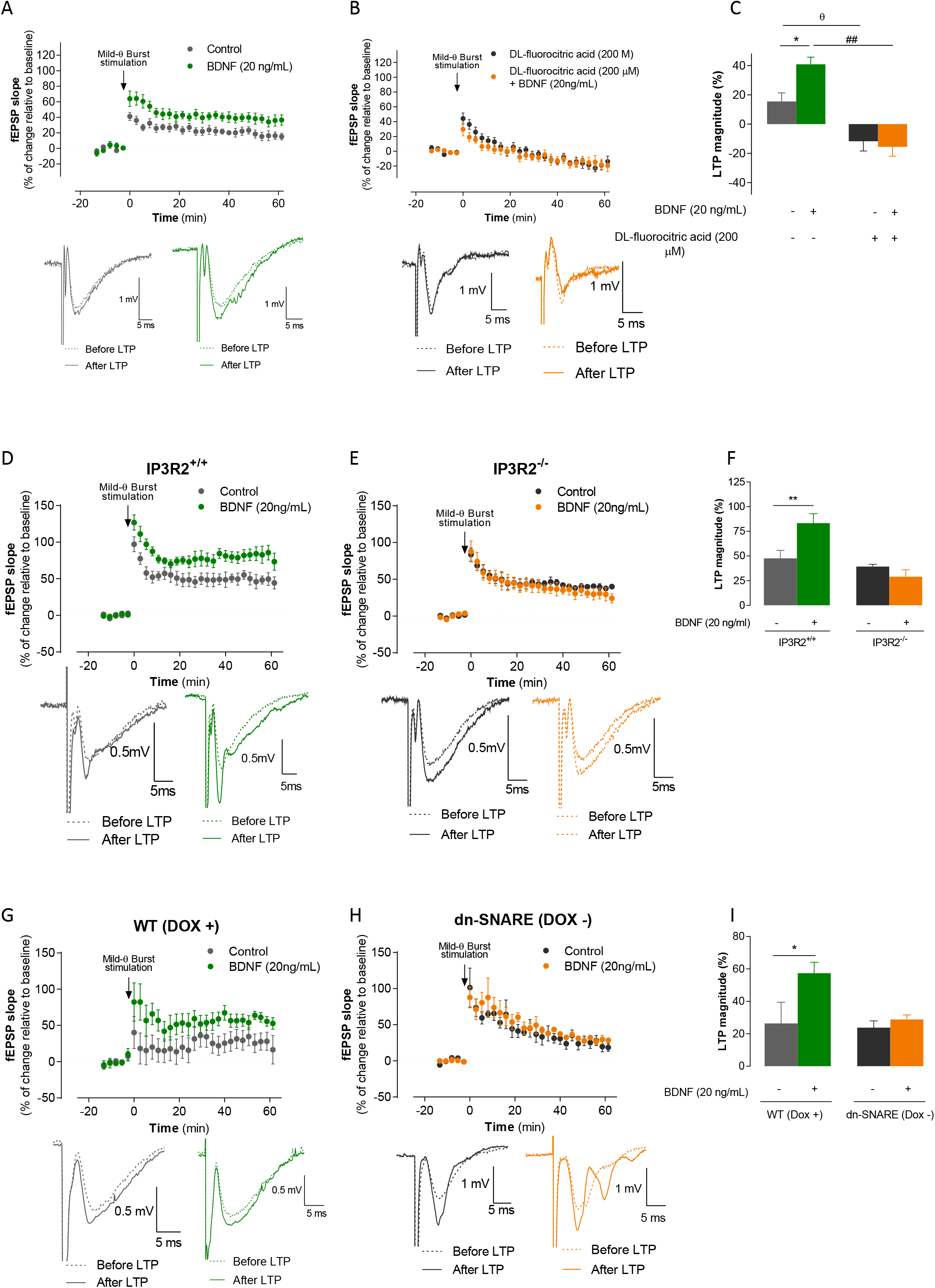
Astrocytic signaling mediates BDNF action under mild-θ-burst induced LTP. In (A) is represented the average time course changes in the fEPSP slope induced by mild θ-burst stimulation in Wistar rats in the absence 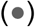 (n=7) or in the presence 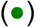 of BDNF (20 ng/mL) (n=6). In (B) is represented the average time course changes in the fEPSP slopes induced by mild θ-burst stimulation in Wistar rats in the presence of FC alone 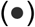 (n=12) or together with BDNF 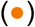 (20 ng/mL) (n=7). In (C) is represented the histogram depict LTP magnitude (change in fEPSP slope at 50-60 min) induced by mild θ-burst stimulation in Wistar rats in the presence or absence of FC and/or BDNF as indicated below each column. (D) represents the time course of averaged normalized changes in fEPSP slope after delivery (arrow) of a mild θ-burst to hippocampal slices from IP3R2^+/+^ in the absence 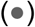 (n= 7) or in the presence 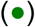 of BDNF (20 ng/mL) (n= 7). (E) Represents the time course of averaged normalized changes in fEPSP slope after delivery (arrow) of a mild θ-burst to hippocampal slices from IP3R2^−/−^ in the absence 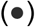 (n= 7) or in the presence 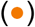 of BDNF (20 ng/mL) (n= 7). (F) corresponds to the histogram depict LTP magnitude (change in fEPSP slope at 50-60 min) induced by mild θ-burst stimulation in the absence or presence of BDNF (20 ng/mL) for each group of animals tested, as indicated below each column. (G) Represents the time course of averaged normalized changes in fEPSP slope after delivery (arrow) of a mild θ-burst to hippocampal slices from WT (Dox +) in the absence 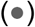 (n= 4) or in the presence 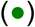 of BDNF (20 ng/mL) (n= 4). (H) Represents the time course of averaged normalized changes in fEPSP slope after delivery (arrow) of a mild θ-burst to hippocampal slices from dn-SNARE (Dox -) in the absence 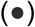 (n= 4) or in the presence 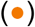 of BDNF (20 ng/mL) (n= 3). (I) corresponds to the histogram depict LTP magnitude (change in fEPSP slope at 50-60 min) induced by mild θ-burst stimulation in the absence or presence of BDNF (20 ng/mL) for each group of animals tested, as indicated below each column. The ordinates in (A), (B), (D), (E), (G) and (H) represent normalized fEPSP slopes where 0% corresponds to the averaged slope recorded for 10 min before θ-burst stimulation and the abscissa represents the time that average begun. Insets bellow each graph correspond to illustrative traces from representative experiments; each trace is the average of eight consecutive responses obtained for each group before (⋯) and 58-60 min after (-) θ-burst stimulation, and is composed of the stimulus artifact, followed by the presynaptic volley and the fEPSP. All values are presented as mean ± standard error of mean (SEM) from *n* independent observations. *p≤ 0.05; **p ≤ 0.01; ^θ^p ≤ 0.01; ^##^p ≤ 0.01 (one-way ANOVA followed by Holm-Sidak’s *post hoc* test for multiple comparisons).

Moreover, in hippocampal slices from IP3R2^+/+^ mice, mild-θ-burst stimulation induced a significant LTP magnitude increase of 47.7±8.01% (n=7, Figure 2E, F). In the same slices, in the presence of BDNF (20ng/mL), the same stimulation paradigm increased the fEPSP slope by 83.3±9.61% (Figure 2D, F; one-way ANOVA following Holm-Sidak’s *post hoc* test; **p=0.006, n=7). Mild-θ-burst stimulation applied in hippocampal slices from IP3R2^−/−^ in the absence of result in an increase of fEPSP of 39.2±2.35%, while in BDNF (20ng/mL) presence LTP magnitude was 29.0±7.07% (Figure 2E, F; one-way ANOVA following Holm-Sidak’s *post hoc* test; n.s, p=0.548, n=7), which corresponds to an impairment of excitatory BDNF effect upon LTP magnitude, indicating that IP3R2-mediated Ca^2+^-signaling controls BDNF effect on hippocampal LTP while using mild-θ-burst stimulation paradigm.

While recording fEPSPs from hippocampal slice of WT (DOX+) mice, a Mild-θ-burst stimulation potentiated the fEPSP slope by 26.4±13.1% (n=4), whereas in the same slices but in the presence of BDNF (20ng/mL) the same induction paradigm enhanced the fEPSP slope by 57.4±6.74% (Figure 2G, I; one-way ANOVA following Holm-Sidak’s *post hoc* test; *p=0.031, n=4) corresponding to a significant effect of BDNF on synaptic plasticity. For dn-SNARE (DOX+) mice we observed a LTP enhancement similar to the WT (DOX+) mice, being the LTP magnitude of 23.2±13.9% (n=4) in the absence of BDNF (20ng/mL) and by 50.4±8.89% (n=4) in its presence (Supp. Fig. 1 D, F; unpaired t-test *p=0.04, n=4). In slices from dn-SNARE (DOX-) mice, the mild-θ-burst stimulation increase LTP magnitude by 23.8±4.19% (n=4) in the absence of BDNF (20ng/mL) and by 28.9±2.7% (n=3) in its presence (Figure 2H, I; one-way ANOVA following Holm-Sidak’s *post hoc* test; n.s, p=0.724, n=3-4), which corresponds to a loss of BDNF action in LTP.

These results show that the compromised astrocytic function, achieved by decreasing astrocytic metabolism, compromising intracellular Ca^2+^-signaling or compromising the release of gliotransmitters through the expression of dn-SNARE domain in astrocytes, leads to the loss of BDNF effect upon LTP under mild stimulation conditions. These results suggest that astrocytes are able to respond differently to diverse intensities of stimulation, and in this way being able to dynamically control BDNF effect on synaptic plasticity.

## Discussion

The main finding of the present study is that astrocytes play an active role in the facilitatory action of BDNF upon hippocampal LTP. In the three studied animal models, pharmacological astrocytic metabolism reduction in Wistar rats, IP3R2−/− mice (which lacks Ca^2+^ signaling in astrocyte) and dn-SNARE transgenic mice (which have astrocytic gliotransmitters release compromised), the effect of BDNF upon LTP was completely abolished, while inducing LTP with a mild-θ-burst stimulation paradigm.

Astrocytes are critical players in diverse aspects of brain function and modulate several cognitive processes, such as learning and memory (Kofuji *et al.*, 2021). Indeed, astrocytes by the Ca^2+^-dependent release of gliotransmitters modulate synaptic plasticity, the physiological basis of learning and memory(Kofuji *et al.*, 2021; Pascual *et al.*, 2005; Henneberger *et al.*, 2010). In the present study we observed an impairment of hippocampal LTP while blocking astrocytes pharmacologically, for both strong and mild- θ-burst stimulation, as well as a decrease in the LTP magnitude in IP3R2−/− mice and dn-SNARE (DOX-) mice, being our results in agreement with literature. Astrocytes modulate LTP, through the release and regulation of different gliotransmitters (Kofuji *et al.*, 2021; Gonçalves-Ribeiro *et al.*, 2019), namely glutamate, D-serine and ATP. Indeed, CA1 hippocampal LTP can be triggered when postsynaptic activity and astrocytic Ca^2+^-dependent glutamate release occur at the same time (Perea *et al.*, 2007); this form of LTP is independent of postsynaptic NMDAR-mediated signaling and requires mGluR activation (Perea *et al.*, 2007). The Ca^2+^-dependent release of D-serine from CA1 hippocampal astrocytes controls NMDAR-dependent plasticity in the excitatory synapses nearby (Henneberger *et al.*, 2010). The extracellular adenosine, derived from astrocytic ATP, regulates synaptic transmission and modulates LTP (Pascual *et al.*, 2005). In spite of some controversy on the physiological relevance of Ca^2+^-dependent exocytosis of gliotransmitters, several studies reported a Ca^2+^-dependent exocytosis of glutamate and ATP from cultured astrocytes (Bal-Price *et al.*, 2002; Petrelli *et al.*, 2016). Recent evidence (Lalo *et al.*, 2014) strongly reinforces the idea that ATP release from astrocytes has a relevant role in astrocytic-to-neuron signaling, namely: 1) the existence of a Cam-dependent mechanism leading to vesicular release of gliotransmitters from neocortical astrocytes, 2) the quantal activation of P2X receptors in neocortical neurons by ATP released from astrocytes, and 3) the glia-driven purinergic modulation of GABAergic transmission that is impaired by astrocytic expression of dn-SNARE (which precludes gliotransmitter release) or deletion of P2X4 receptors.

Brain-derived neurotrophic factor is a key neurotrophin which plays a critical modulatory action upon neuronal survival, dendritic growth and plasticity (Gottschalk *et al.*, 1999; Carvalho *et al.*, 2008; Minichiello, 2009; Graciano Leal *et al.*, 2015). In the present work we also observed a potentiation of LTP magnitude in the presence of BDNF, nevertheless the effect was only observed when inducing LTP with a strong θ-burst stimulation, and it was not observed while using a mild θ-burst stimulation. Mainly, BDNF mediated is actions through the activation of TrkB-FL receptors, which are coupled to three different signaling pathways: (i) the phosphatidylinositol-3-kinase (PI3K)/Akt pathway, (ii) the Ras/MAPK pathway, and (iii) the PLCγ pathway (Gottschalk *et al.*, 1999). Working together, these pathways underlie important cognitive processes, including learning and memory formation. In fact, it is known that BDNF leads to an increase in LTP magnitude, by its binding to TrkB-FL receptors, both at a presynaptic and postsynaptic level (Figurov *et al.*, 1996). Several studies reported that BNDF effect on LTP is depend of adenosine A2A receptor (Fontinha *et al.*, 2008), being the excitatory action of BDNF on hippocampal LTP fully dependent on the recruitment and activation of A2AR, with the following activation of cAMP/protein kinase A (PKA) signaling cascade (Fontinha *et al.*, 2008). Thus, adenosine released by astrocytes may account for the BDNF excitatory effect on LTP only for mild θ-burst stimulation, while under strong θ-burst stimulation the necessary adenosine for BDNF effect is provided by astrocytes. Thus we are showing that astrocytes can detect and decode differences in neuronal activity and respond to it by the release of gliotransmitters. Since BNDF action upon LTP is depend of A2A receptors is possible that the gliotransmitters involved in this process is ATP/adenosine, and this remains to be evaluated. In this study, the use of different stimulation protocols allows to decipher the distinct astrocytic actions upon different stimulation contexts, where for mild-θ-burst induced hippocampal LTP, astrocytic gliotransmitters would be found to be essential for the facilitatory BDNF action. However, for a strong-θ-burst paradigm, astrocytic gliotransmitter were not necessary for BDNF action upon LTP. Therefore, in agreement with previous literature (Covelo *et al.*, 2018), the present study presents evidences that astrocytes can detect and decode neuronal activity and respond distinctly to it.

Hence, this data shows for the first time that astrocytes play an active role in the facilitatory action of BDNF upon LTP and suggests that they do so by being a source of the gliotransmitters.

## Acknowledgments

This work was supported by project funding from Fundação para a Ciência e para a Tecnologia (FCT) to SHV (PTDC/BTM-SAL/32147/2017) and AMS (PTDC/MED-FAR/30933/2017). This project has received funding from H2020-WIDESPREAD-05-2017-Twinning (EpiEpinet) under grant agreement No. 952455. AAM (PD/BD/114278/2016), JG-R (PD/BD/150342/2019), and SP (SFRH/BD/147277/2019) are supported by PhD fellowships from FCT.

We thank Regeneron Pharmaceuticals for the gift of brain-derived neurotrophic factor.

The authors are grateful to Prof. Philip Haydon, Prof. Ju Chen and Prof João Oliveira for sharing the dn-SNARE and IP3R2 mice lines.

---

**Supplementary Figure 1.**
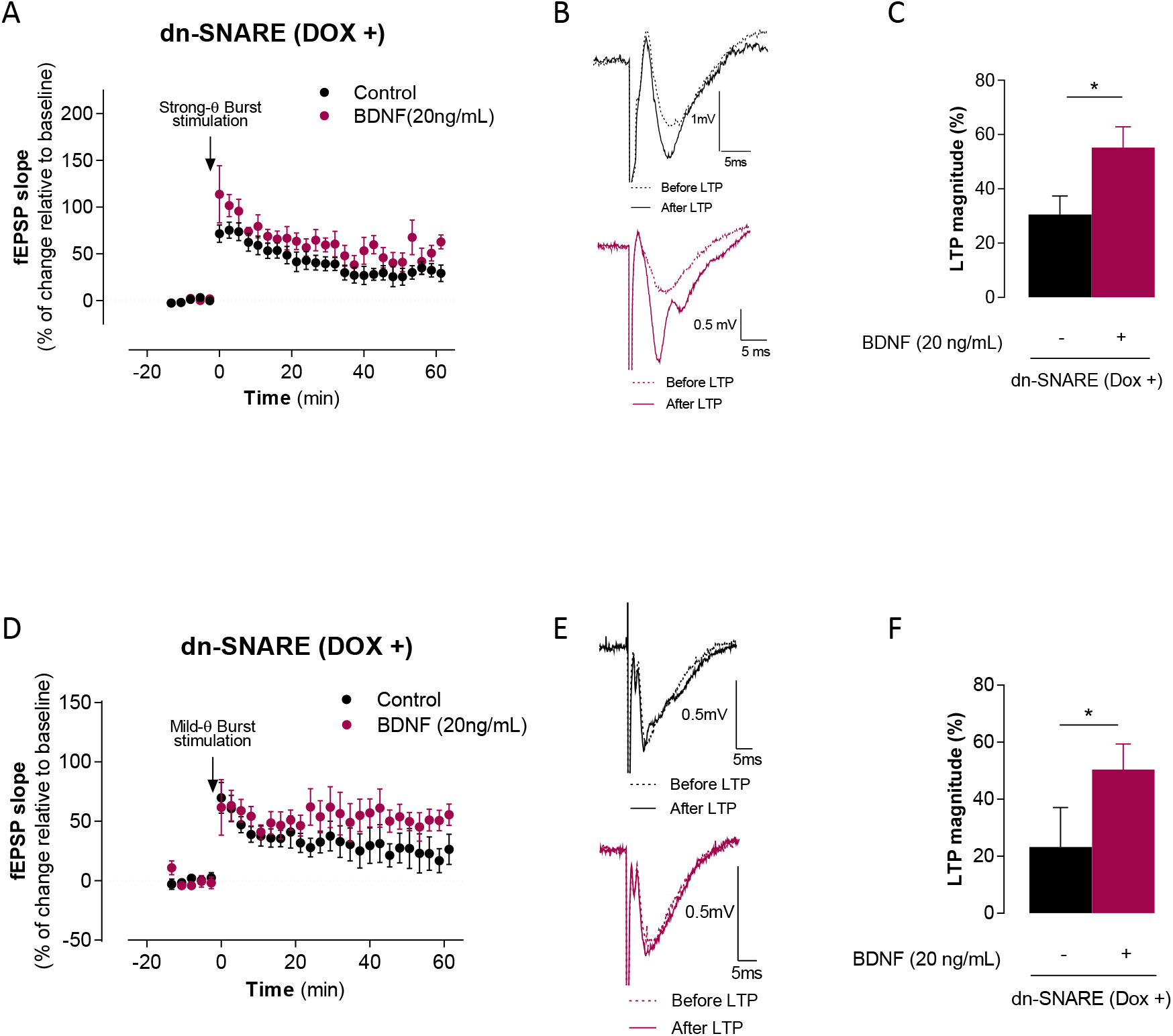
BDNF effect over LTP is dependent on the astrocytic-release of gliotransmitters via exocytosis. (A) Represents the time course of averaged normalized changes in fEPSP slope after delivery (arrow) of a strong θ-burst to hippocampal slices from dn-SNARE (Dox +) in the absence 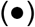 (n=5) or in the presence 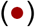 of BDNF (20 ng/mL) (n= 3). (B) corresponds to illustrative traces from representative experiments; each trace is the average of eight consecutive responses obtained for each group before (⋯) and 58-60 min after (–) θ-burst stimulation, and is composed of the stimulus artifact, followed by the presynaptic volley and the fEPSP. (C) corresponds to the histogram depict LTP magnitude (change in fEPSP slope at 50-60 min) induced by strong θ-burst stimulation in hippocampal slices from dn-SNARE (Dox +) mice in the presence or absence of BDNF (20 ng/mL), as indicated below each column. (D) Represents the time course of averaged normalized changes in fEPSP slope after delivery (arrow) of a mild θ-burst to hippocampal slices from dn-SNARE (Dox +) in the absence 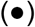 (n=4) or in the presence 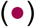 of BDNF (20 ng/mL) (n= 4). (B) correspond to illustrative traces from representative experiments; each trace is the average of eight consecutive responses obtained for each group before (⋯) and 58-60 min after (–) θ-burst stimulation, and is composed of the stimulus artifact, followed by the presynaptic volley and the fEPSP. (C) corresponds to the histogram depict LTP magnitude (change in fEPSP slope at 50-60 min) induced by mild θ-burst stimulation in hippocampal slices from dn-SNARE (Dox +) mice in the presence or absence of BDNF (20 ng/mL), as indicated below each column. The ordinates in (A) and (D) represent normalized fEPSP slopes where 0% corresponds to the averaged slope recorded for 10 min before θ-burst stimulation and the abscissa represents the time that average begun. Insets bellow each graph correspond to illustrative traces from representative experiments; each trace is the average of eight consecutive responses obtained for each group before (⋯) and 58-60 min after (–) θ-burst stimulation, and is composed of the stimulus artifact, followed by the presynaptic volley and the fEPSP. All values are presented as mean ± standard error of mean (SEM) from *n* independent observations. Statistical significance was assessed by unpaired t-test between control 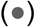 and BDNF (20 ng/mL) 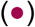 conditions. p> 0.05; *p≤ 0.05.

## References

Albensi, Benedict C., Oliver, Derek R., Toupin, Justin, and Odero, Gary. (2007). Electrical Stimulation Protocols for Hippocampal Synaptic Plasticity and Neuronal Hyper-Excitability: Are They Effective or Relevant? Experimental Neurology, 204, 1–13.

Araque, Alfonso, Parpura, Vladimir, Sanzgiri, Rita P., and Haydon, Philip G. (1999). Tripartite Synapses: Glia, the Unacknowledged Partner. Trends in Neurosciences,.

Bal-Price, Anna, Moneer, Zahid, and Brown, Guy C. (2002). Nitric Oxide Induces Rapid, Calcium-Dependent Release of Vesicular Glutamate and ATP from Cultured Rat Astrocytes. Glia, 40, 312–23.

Berg-Johnsen, J., Paulsen, R. E., Fonnum, F., and Langmoen, I. A. (1993). Changes in Evoked Potentials and Amino Acid Content during Fluorocitrate Action Studied in Rat Hippocampal Cortex. Experimental Brain Research, 96, 241–46.

Bliss, T. V. P., and Collingridge, G. L. (1993). A Synaptic Model of Memory: Long-Term Potentiation in the Hippocampus. Nature,.

Carvalho, a L, Caldeira, M V, Santos, S D, and Duarte, C B. (2008). Role of the Brain-Derived Neurotrophic Factor at Glutamatergic Synapses. British Journal of Pharmacology, 153 Suppl, S310–24.

Covelo, Ana, and Araque, Alfonso. (2018). Neuronal Activity Determines Distinct Gliotransmitter Release from a Single Astrocyte. eLife, 7, 1–19.

Figurov, Alexander, Pozzo-Miller, Lucas D., Olafsson, Petur, Wang, Ti, and Lu, Bai. (1996). Regulation of Synaptic Responses to High-Frequency Stimulation and LTP by Neurotrophins in the Hippocampus. Nature, 381, 706–9.

Fontinha, B M, Diógenes, M J, Ribeiro, J a, and Sebastião, a M. (2008). Enhancement of Long-Term Potentiation by Brain-Derived Neurotrophic Factor Requires Adenosine A2A Receptor Activation by Endogenous Adenosine. Neuropharmacology, 54, 924–33.

Gonçalves-Ribeiro, Joana, Pina, Carolina Campos, Sebastião, Ana Maria, and Vaz, Sandra Henriques. (2019). Glutamate Transporters in Hippocampal LTD/LTP: Not Just Prevention of Excitotoxicity. Frontiers in Cellular Neuroscience,.

Gottschalk, W.A., Jiang, H., Tartaglia, N., Feng, L., Figurov, A., and Lu, B. (1999). Signaling Mechanisms Mediating BDNF Modulation of Synaptic Plasticity in the Hippocampus. Learn.Mem., 6, 243–56.

Hassel, Bjørnar, Sonnewald, Ursula, Unsgård, Geirmund, and Fonnum, Frode. (1994). NMR Spectroscopy of Cultured Astrocytes: Effects of Glutamine and the Gliotoxin Fluorocitrate. Journal of Neurochemistry, 62, 2187–94.

Henneberger, Christian, Papouin, Thomas, Oliet, Stéphane H R, and Rusakov, Dmitri a. (2010). Long-Term Potentiation Depends on Release of D-Serine from Astrocytes. Nature, 463, 232–36.

Jerónimo-Santos, André, Vaz, Sandra Henriques, Parreira, Sara, Rapaz-Lérias, Sofia, Caetano, António P, Buée-Scherrer, Valérie, Castrén, Eero, et al. (2014). Dysregulation of TrkB Receptors and BDNF Function by Amyloid-β Peptide Is Mediated by Calpain. Cerebral Cortex (New York, N.Y.: 1991),.

Kofuji, Paulo, and Araque, Alfonso. (2021). Astrocytes and Behavior. Annual Review of Neuroscience, 44,.

Korte, Martin, Carroll, Patrick, Wolf, Eckhard, Brem, Gottfried, Thoenen, Hans, and Bonhoeffer, Tobias. (1995). Hippocampal Long-Term Potentiation Is Impaired in Mice Lacking Brain-Derived Neurotrophic Factor. Proceedings of the National Academy of Sciences of the United States of America, 92, 8856–60.

Lalo, Ulyana, Palygin, Oleg, Rasooli-Nejad, Seyed, Andrew, Jemma, Haydon, Philip G., and Pankratov, Yuriy. (2014). Exocytosis of ATP From Astrocytes Modulates Phasic and Tonic Inhibition in the Neocortex. PLoS Biology, 12,.

Largo, Carlota, Cuevas, Pedro, Somjen, George G., Martín Del Río, Rafael, and Herreras, Oscar. (1996). The Effect of Depressing Glial Function in Rat Brain in Situ on Ion Homeostasis, Synaptic Transmission, and Neuron Survival. Journal of Neuroscience, 16, 1219–29.

Leal, G., Bramham, C. R., and Duarte, C. B. (2017). BDNF and Hippocampal Synaptic Plasticity. In Vitamins and Hormones, 104:153–95.

Leal, Graciano, Afonso, Pedro M., Salazar, Ivan L., and Duarte, Carlos B. (2015). Regulation of Hippocampal Synaptic Plasticity by BDNF. Brain Research, 1621, 82–101.

Li, Xiaodong, Zima, Aleksey V., Sheikh, Farah, Blatter, Lothar a., and Chen, Ju. (2005). Endothelin-1-Induced Arrhythmogenic Ca2+ Signaling Is Abolished in Atrial Myocytes of Inositol-1,4,5-trisphosphate(IP3)-Receptor Type 2-Deficient Mice. Circulation Research, 96, 1274–81.

Minichiello, Liliana. (2009). TrkB Signalling Pathways in LTP and Learning. Nature Reviews. Neuroscience, 10, 850–60.

Minichiello, Liliana, Calella, Anna Maria, Medina, Diego L., Bonhoeffer, Tobias, Klein, Rüdiger, and Korte, Martin. (2002). Mechanism of TrkB-Mediated Hippocampal Long-Term Potentiation. Neuron, 36, 121–37.

Pascual, Olivier, Casper, Kristen B, Kubera, Cathryn, Zhang, Jing, Revilla-Sanchez, Raquel, Sul, Jai-Yoon, Takano, Hajime, Moss, Stephen J, McCarthy, Ken, and Haydon, Philip G. (2005). Astrocytic Purinergic Signaling Coordinates Synaptic Networks. Science (New York, N.Y.), 310, 113–16.

Patterson, Susan L., Abel, Ted, Deuel, Thomas A.S., Martin, Kelsey C., Rose, Jack C., and Kandel, Eric R. (1996). Recombinant BDNF Rescues Deficits in Basal Synaptic Transmission and Hippocampal LTP in BDNF Knockout Mice. Neuron, 16, 1137–45.

Paulsen, R. E., Contestabile, A., Villani, L., and Fonnum, F. (1987). An In Vivo Model for Studying Function of Brain Tissue Temporarily Devoid of Glial Cell Metabolism: The Use of Fluorocitrate. Journal of Neurochemistry, 48, 1377–85.

Perea, Gertrudis, and Araque, Alfonso. (2007). Astrocytes Potentiate Transmitter Release at Single Hippocampal Synapses. Science (New York, N.Y.), 317, 1083–86.

Petrelli, Francesco, and Bezzi, Paola. (2016). Novel Insights into Gliotransmitters. Current Opinion in Pharmacology,.

Pins, Benoit De, Cifuentes-Díaz, Carmen, Thamila Farah, Amel, López-Molina, Laura, Montalban, Enrica, Sancho-Balsells, Anna, López, Ana, et al. (2019). Conditional BDNF Delivery from Astrocytes Rescues Memory Deficits, Spine Density, and Synaptic Properties in the 5xFAD Mouse Model of Alzheimer Disease. Journal of Neuroscience, 39, 2441–58.

Rei, N., Rombo, D. M., Ferreira, M. F., Baqi, Y., Müller, C. E., Ribeiro, J. A., Sebastião, A. M., and Vaz, S. H. (2020). Hippocampal Synaptic Dysfunction in the SOD1G93A Mouse Model of Amyotrophic Lateral Sclerosis: Reversal by Adenosine A2AR Blockade. Neuropharmacology, 171,.

Rose, Christine R, Blum, Robert, Pichler, Bruno, Lepier, Alexandra, Kafitz, Karl W, and Konnerth, Arthur. (2003). Truncated TrkB-T1 Mediates Neurotrophin-Evoked Calcium Signalling in Glia Cells. Nature, 426, 74–78.

Sardinha, Vanessa Morais, Guerra-Gomes, Sónia, Caetano, Inês, Tavares, Gabriela, Martins, Manuella, Reis, Joana Santos, Correia, Joana Sofia, et al. (2017). Astrocytic Signaling Supports Hippocampal–prefrontal Theta Synchronization and Cognitive Function. GLIA, 65, 1944–60.

Sebastião, Ana M, Assaife-Lopes, Natália, Diógenes, Maria J, Vaz, Sandra H, and Ribeiro, Joaquim a. (2011). Modulation of Brain-Derived Neurotrophic Factor (BDNF) Actions in the Nervous System by Adenosine A(2A) Receptors and the Role of Lipid Rafts. Biochimica et Biophysica Acta, 1808, 1340–49.

Sultan, Sébastien, Li, Liyi, Moss, Jonathan, Petrelli, Francesco, Cassé, Frédéric, Gebara, Elias, Lopatar, Jan, et al. (2015). Synaptic Integration of Adult-Born Hippocampal Neurons Is Locally Controlled by Astrocytes. Neuron, 88, 957–72.

Swanson, Raymond A., and Graham, Steven H. (1994). Fluorocitrate and Fluoroacetate Effects on Astrocyte Metabolism in Vitro. Brain Research, 664, 94–100.

Todd, Keith J., Auld, Daniel S., and Robitaille, Richard. (2007). Neurotrophins Modulate Neuron-Glia Interactions at a Vertebrate Synapse. European Journal of Neuroscience, 25, 1287–96.

Xu, Baoji, Gottschalk, Wolfram, Chow, Ana, Wilson, Rachel I., Schnell, Eric, Zang, Keling, Wang, Denan, Nicoll, Roger A., Lu, Bai, and Reichardt, Louis F. (2000). The Role of Brain-Derived Neurotrophic Factor Receptors in the Mature Hippocampus: Modulation of Long-Term Potentiation through a Presynaptic Mechanism Involving trkB. Journal of Neuroscience, 20, 6888–97.

